# High specificity of widely used phospho-tau antibodies validated using a quantitative whole-cell based assay

**DOI:** 10.1101/612911

**Authors:** Dan Li, Yong Ku Cho

**Affiliations:** Departments of Biomedical Engineering, University of Connecticut, Storrs, CT 06269; Departments of Chemical and Biomolecular Engineering, University of Connecticut, Storrs, CT 06269; Institute for Systems Genomics, University of Connecticut, Storrs, CT 06269; Connecticut Institute for the Brain and Cognitive Sciences, University of Connecticut, Storrs, CT 06269

**Author notes:** To whom correspondence should be addressed: Yong Ku Cho: Departments of Chemical and Biomolecular Engineering and Biomedical Engineering, Institute for Systems Genomics, University of Connecticut, Storrs CT 06269.

**Keywords:** Antibody, Tau protein, protein phosphorylation, phospho-tau, antibody specificity, post-translational modification

## Abstract

Antibodies raised against defined phosphorylation sites of the microtubule-associated protein tau are widely used in scientific research and being applied in clinical assays. However, recent studies have revealed an alarming degree of non-specific binding found in these antibodies. In order to quantify and compare the specificity phospho-tau antibodies and other post-translational modification site-specific antibodies in general, a measure of specificity is urgently needed. Here we report a robust flow cytometry assay using human embryonic kidney (HEK) cells that enables the determination of a specificity parameter termed Φ, which measures the fraction of non-specific signal in antibody binding. We validate our assay using anti-tau antibodies with known specificity profiles, and apply it to measure the specificity of 7 widely used phospho-tau antibodies (AT270, AT8, AT100, AT180, PHF-6, TG-3, and PHF-1) among others. We successfully determined the Φ values for all antibodies except AT100, which did not show detectable binding in our assay. Our results show that antibodies AT8, AT180, PHF-6, TG-3, and PHF-1 have Φ values near 1, which indicates no detectable non-specific binding. AT270 showed Φ value around 0.8, meaning that approximately 20% of the binding signal originates from non-specific binding. Further analyses using immunocytochemistry and western blotting confirmed the presence of non-specific binding of AT270 to non-tau proteins found in HEK cells and the mouse hippocampus. We anticipate that the quantitative approach and parameter introduced here will be widely adopted as a standard for reporting the specificity for phospho-tau antibodies, and potentially for post-translational modification targeting antibodies in general.

## Introduction

Site-specific phosphorylation of the microtubule-associated protein tau is a well-defined molecular signature linked to normal tau function as well as pathology in neurodegeneration. Antibodies binding to phosphorylated tau (phospho-tau) are extensively used in research and being adopted in clinical diagnosis (Chai *et al.* 2011; Boutajangout *et al.* 2011; Kayed 2010; Asuni *et al.* 2007; Boutajangout *et al.* 2010; Lewis *et al.* 2000). According to antibody databases (Antibodypedia and Alzforum), thousands of anti-tau antibodies are currently available, including >180 unique phospho-tau antibodies. However, few validation data have been published (Petry *et al.* 2014; Ercan *et al.* 2017; Mercken *et al.* 1992), which indicated an alarming degree of non-specific binding frequently observed in phospho-tau antibodies. To prevent waste of resource and irreproducibility of results, a standard measure of specificity, equivalent to dissociation constants (*K*_d_) for measuring affinity, is urgently needed. Ideally, this parameter should enable users to compare the specificity, identify the source of non-specificity, and clearly understand what is measured.

Existing specificity characterization methods relied on immunoblotting approaches using either synthetic peptides, cell lysates or extracted paired helical filament (PHF) tau (Petry *et al.* 2014; Ercan *et al.* 2017; Mercken *et al.* 1992). Synthetic peptides allow binding measurement to exactly defined phosphorylation sites, but is prohibitively costly to cover all phospho-sites across the full-length tau. Cell lysates allow detection of non-specific binding to non-tau proteins (for example by using cell lysates from a tau knockout mouse) (Petry *et al.* 2014), but cannot measure cross-reactivity to non-modified tau or other phosphorylation sites. Extracted PHF-tau from AD brain homogenates allows characterizing antibodies that do not recognize normal human adult tau and measuring the cross-reactivity to nonphospho-tau by treatment with phosphatase, but cannot capture the phospho-site specificity (Mercken *et al.* 1992). Steinhilb and colleagues have generated transgenic flies expressing tau with alanine point mutations at several serine-proline and threonine-proline motifs of human tau (Steinhilb *et al.* 2007). In this study, they used fly head extracts from these transgenic lines to perform western blotting using phospho-tau antibodies. In principle, this approach may be applied for validating phospho-tau antibodies. However, all of these approaches are not ideal for quantitative measurement due to frequent signal saturation in immunoblotting experiments (Janes 2015). To fill this critical need, we introduce a parameter termed Φ (phi) that quantifies specificity of phospho-tau antibodies and establish a generalizable, robust assay to measure it.

The approach leverages flow cytometry for quantitative, multi-dimensional analysis of small changes in antibody binding to different cell populations. In a typical immunodetection experiment, signal from specific and non-specific binding cannot be distinguished. We sought to enable this distinction within a single sample, by generating two populations of cells with distinct fluorescence reporters. In one population of cells, the target protein (i.e., tau) is expressed as a fusion to the enhanced green fluorescent protein (EGFP). In another population, tau containing a site-specific alanine mutation is expressed as a fusion to a near infra-red fluorescent protein (iRFP). The two populations of cells are mixed, and labeled with a phospho-specific antibody (**Fig. 1**). For a phospho-site specific antibody, any binding to iRFP-positive cells is non-specific binding, enabling the measurement of the fraction of non-specific signal among all antibody binding signals. In order to ensure the measurement of cross-reactivity to non-phosphorylated tau, the experiment is repeated by replacing alanine mutant expressing cells with tau expressing cells (iRFP fused to wild-type tau) treated with the lambda protein phosphatase (λPP). Analogous to the alanine mutation assay, binding to iRFP-positive cells is identified as non-specific signal. Based on these measurements, the fraction of non-specific signal is calculated, which is subtracted from 1 to obtain Φ. Therefore, Φ indicates the fraction of specific signal.

**Figure 1.**
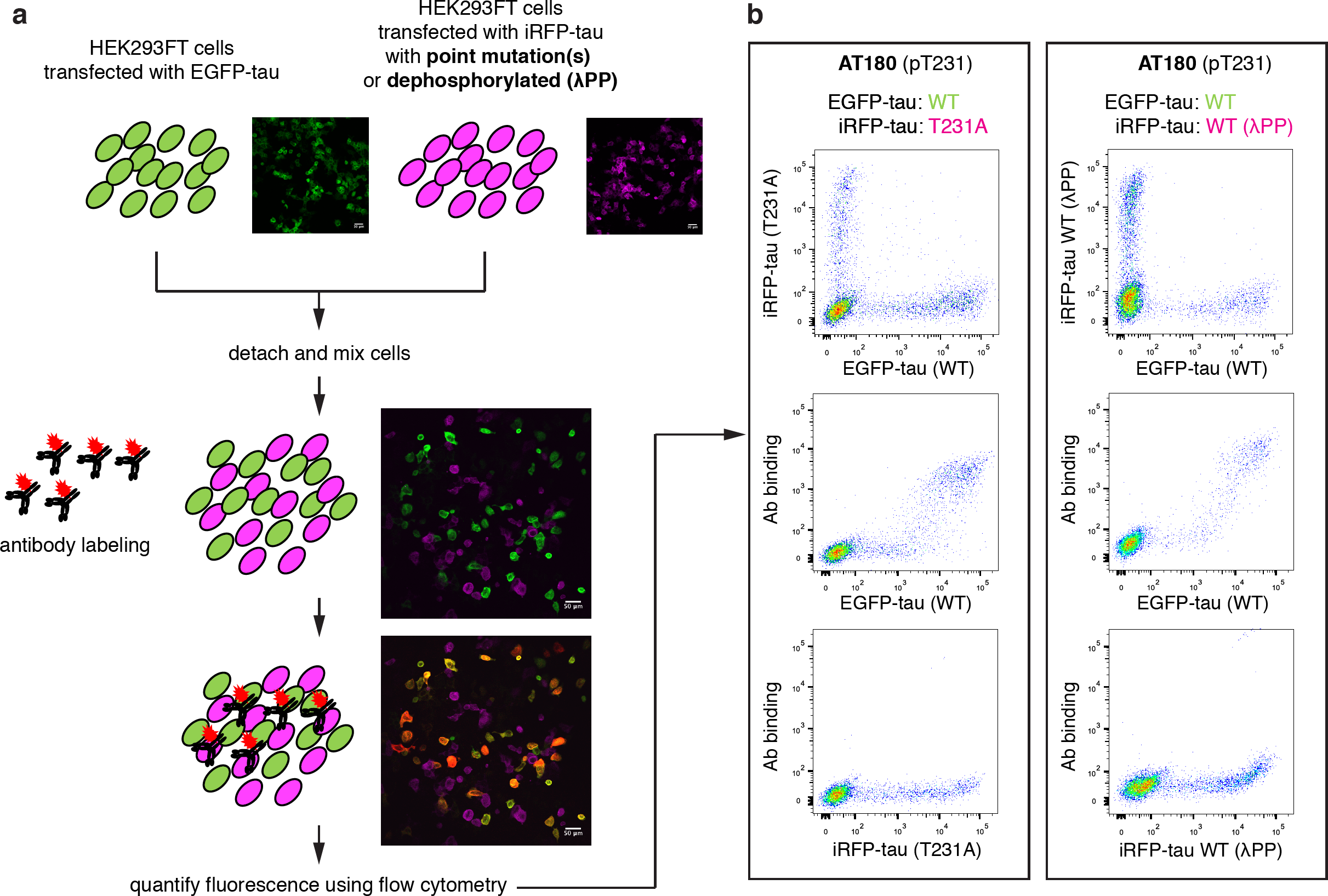
A ratiometric assay to quantify the specificity of phospho-tau antibodies. **a**, Schematic of phospho-tau antibody specificity quantification assay using HEK293FT. HEK293FT cells were transfected with either EGFP-tau WT (green) or iRFP-tau with alanine mutation(s) at the target site (magenta), in separate wells. The cells were then mixed and incubated with phospho-tau antibodies. The antibody binding was detected using an Alexa Flour 594 conjugated secondary antibody (red). In parallel, HEK293FT cells were transfected with either EGFP-tau WT (green) or iRFP-tau WT (magenta), in separate wells. iRFP-tau WT cells were treated with lambda phosphatase (λPP) to de-phosphorylate tau, then mixed with EGFP-tau WT cells. Scale bars, 50 μm. **b**, Representative flow cytometry results for AT180 showing the alanine mutant experiment (left) and de-phosphorylation experiment (right).

Using this measurement, we measured the specificity Φ of 7 commonly used phospho-tau antibodies (AT270, AT8, AT100, AT180, PHF-6, TG-3, and PHF-1), in addition to others. Our results show that 5 antibodies (AT8, AT180, PHF-6, TG-3, and PHF-1) have Φ values near 1, indicating nearly perfect specificity. Antibodies AT270, AT100 showed weak but detectable non-specific binding to non-tau proteins in cells, with lower Φ values. This resulted in background fluorescence in flow cytometry and immunocytochemistry experiments. The non-specific binding also resulted in false positive detection in western blots, evidenced by additional bands appearing in cell lysates that do not contain human tau. To our knowledge, this is the first quantitative measurement of antibody specificity that enables specificity comparison with information on the source of non-specificity.

## Methods

### Cloning of recombinant constructs of tau and mutant tau

The following plasmids were obtained from Addgene: a mammalian expression vector pRK5 encoding EGFP tagged wild-type (WT) human tau (pRK5-EGFP-tau, # 46904, RRID: Addgene_46904), which contains four C-terminal repeat regions and lacks the N-terminal sequences (0N4R), a plasmid encoding iRFP (piRFP670-N1, # 44457, RRID: Addgene_45457), and a mammalian expression vector pcDNA3 encoding hemagglutinin tagged constitutively active glycogen synthase kinase-3β (GSK-3β) (S9A) (pcDNA3-GSK3β, # 14754, RRID: Addgene_14754). The EGFP in pRK5 vector was first replaced by iRFP to construct an iRFP tagged WT tau by golden gate assembly (pRK5-iRFP-tau) (Engler *et al.* 2008). This assembly was performed using a Type IIs restriction enzyme BsaI (New England Biolabs # R3535) and T4 DNA ligase (New England Biolabs # M0202) (see **Supp. Table 1** for primers used). Seven alanine mutant variants were generated in iRFP tagged tau to preclude phosphorylation. Alanine mutations were introduced to either Thr181, Ser202, Thr212/Ser214 (double alanine mutant), Thr231, Ser262, Ser396/Ser404 (double alanine mutant), or Ser404. For this, site-directed mutagenesis was conducted using overlap extension PCR (Pavoor *et al.* 2009) using the Phusion DNA polymerase (New England Biolabs # M0530) (see **Supp. Table 2**. for primers used). Residue numbering of tau is according to the longest adult human brain tau isoform.

**Table 1.**

| Name | Target Phospho-site | Host/ Isotype | Class | Source | Dilution | Working conc. | $\Phi_{ala}$ | $\Phi_{\lambda PP}$ | Reference |
| --- | --- | --- | --- | --- | --- | --- | --- | --- | --- |
| **AT270** | pT181 | Mouse/IgG1, $\kappa$ | Monoclonal | Invitrogen, # MN1050, RRID: AB_223651 | 1:100 | 2 μg/mL | 0.80±0.06 | 0.99±0.01 | (Goedert *et al.* 1994; Caccamo *et al.* 2010) |
| **AT8** | pS202/pT205 | Mouse/IgG1 | Monoclonal | Invitrogen, # MN1020, RRID: AB_223647 | 1:100 | 2 μg/mL | 0.95±0.07 | 0.95±0.05 | (Goedert *et al.* 1995; Gotz *et al.* 1995; Gotz *et al.* 2001; Yanamandra *et al.* 2013) |
| **AT100** | pT212/pS214 | Mouse/IgG1, $\kappa$ | Monoclonal | Invitrogen, # MN1060, RRID: AB_223652 | 1:100 | 2 μg/mL | N/A | N/A | (Mercken *et al.* 1992; Gotz *et al.* 2001; Jackson *et al.* 2002) |
| **AT180** | pT231 | Mouse/IgG1, $\kappa$ | Monoclonal | Invitrogen, # MN1040, RRID: AB_223649 | 1:100 | 2 μg/mL | 0.99±0.01 | 0.98±0.01 | (Goedert *et al.* 1994; Gotz *et al.* 2001; Nakamura *et al.* 2012) |
| **PHF6** | pT231 | Mouse/IgG1 | Monoclonal | Invitrogen, # 35-5200, RRID: AB_2533210 | 1:250 | 2 μg/mL | 0.98±0.01 | 0.97±0.00 | (Hoffmann *et al.* 1997; Sankaranarayanan *et al.* 2015) |
| **1H6L6** | pT231 | Rabbit/IgG | Monoclonal | Invitrogen, # 701056, RRID: AB_2532360 | 1:250 | 2 μg/mL | 1.00±0.00 | 0.99±0.01 | (Dengler-Crish *et al.* 2017) |
| **TG-3** | pT231 | Mouse/IgM | Monoclonal | Peter Davies RRID: AB_2716726 | 1:100 | N/A | 0.96±0.01 | 0.95±0.06 | (Dickson *et al.* 1995; Jicha *et al.* 1997; Gotz *et al.* 2001) |
| **pS262** | pS262 | Rabbit/IgG | Polyclonal | Abcam, # ab131354, RRID: AB_11156689 | 1:500 | 2 μg/mL | 0.32±0.02 | 0.37±0.01 | |
| **PHF1** | pS396/pS404 | Mouse/IgG | Monoclonal | Peter Davies RRID: AB_2315150 | 1:100 | N/A | 0.98±0.00 | 0.98±0.03 | (Greenberg *et al.* 1992; Otvos *et al.* 1994; Gotz *et al.* 1995) |
| **pS404** | pS404 | Rabbit/IgG | Monoclonal | Abcam, # ab92676, RRID: AB_10561457 | 1:200 | 0.1 μg/mL | 1.00±0.00 | 0.99±0.00 | |

### Cell culture and transfection

The human embryonic kidney (HEK) 293FT cells a high transfection efficiency isolate of the HEK293T cells were purchased from ThermoFisher (# R70007), and used without further authentication. The cell line is not listed as a commonly misidentified cell line by the International Cell Line Authentication Committee. HEK293FT cells were grown in DMEM supplemented with 10 % fetal bovine serum. The cells were frozen at early passage and used for less than 2 months in continuous culture. HEK293FT cells were seeded in 24-well plate in the same number 2 days before plasmid transfection to reach 70 % confluency. The cells were then transfected with plasmid using the Lipofectamine 3000 reagent (Invitrogen # L3000001) according to the manufacturer’s instructions.

### Tau antibodies

10 phospho-tau antibodies raised against 7 phosphoepitopes in tau were validated for their specificity (**Table 1**). The following monoclonal antibodies were commercially available: AT270 (pThr181), AT8 (pSer202/pThr205), AT100 (pSer212/pSer214), AT180 (pThr231), PHF-6 (pThr231), 1H6L6 (pThr231), and anti-tau (pSer404). The following monoclonal antibodies were a gift of Dr. Peter Davies: TG-3 (pThr231 and conformation-specific) and PHF-1 (pSer396/pSer404). One purified rabbit polyclonal antibody anti-tau (pSer262) was purchased from Abcam. Tau-5 (Tau210-241, Invitrogen # AHB0042, AB_2536235) is a monoclonal antibody recognizing phosphorylated and non-phosphorylated tau isoforms. Purified single-chain antibody variable fragment (scFv) 3.24 targeting pThr231 was reported previously (Li *et al.* 2018).

### Quantitation of antibody specificity

Here we define a generally applicable parameter to quantify antibody specificity, based on general discussions regarding multi-partner protein interaction by Hall and Daugherty (Hall & Daugherty 2009). If an antibody *Ab* is capable of binding to *n* different partners *X*_*1*_, *X*_*2*_, *… X*_*n*_ as described by the following reactions,

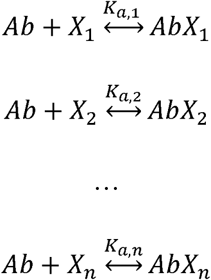

the specificity of the interaction between *Ab* and *X*_*i*_ can be defined as Φ_*i*_ where:

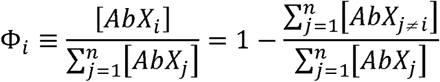

From this definition, 0< *Φ*_*i*_ < 1, where 1 indicates perfect specificity (*Ab* only reacts with its intended antigen *X*_*i*_), and 0 indicates no interaction with *X*_*i*_. We used the following formula to quantify the specificity of phospho-tau antibodies:

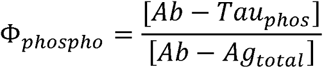

where *Tau*_*phos*_ indicates phosphorylated epitope sequences of tau at a defined site, and *Ag*_*total*_ indicates overall potential partners including *Tau*_*phos*_. In our assay, we determined the specificity using a three-color labeling scheme, where the mixture of HEK293FT cells transiently expressing EGFP tagged WT human tau and iRFP tagged alanine mutant tau was incubated with a fluorescence-labeled antibody. In this case, the overall specificity Φ_*ala*_ can be obtained by measuring the following parameter:

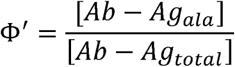

where *Ag*_*ala*_ indicates antigen with an alanine mutation at the target phosphorylation site. Therefore, Φ*′* in this case indicates the fraction of total binding signal to the alanine mutant (non-specific binding signal from cells expressing the alanine mutant). Φ_*ala*_ is defined as:

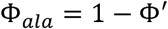

Therefore, Φ_*ala*_ value of 1 indicates no cross-reactivity to the alanine mutant. Experimentally, Φ*′* was determined by measuring the fluorescence intensity of antibody labeling from each cell in flow cytometry experiments (detailed description on how Φ values were obtained is below). For example,

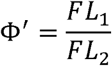

where *FL*_*1*_ and *FL*_*2*_ are fluorescence intensity of antibody labeling from HEK293FT cells transiently expressing alanine mutant tau and WT tau, respectively. Fluorescence intensity was calculated by subtracting the background fluorescence measured on the same day from a control antibody with matching isotype. Analogously, Φ_λPP_ is defined by replacing *Ag*_*ala*_ with lambda protein phosphatase (λPP)-treated antigen (*Ag*_λPP_). Φ_λPP_ value of 1 indicates no cross-reactivity to the de-phosphorylated tau.

### Measurement of Φ: validation of phospho-tau antibody specificity

To measure Φ_*ala*_ of phospho-tau antibodies, the HEK293FT cells were co-transfected with plasmids pRK5-EGFP-tau WT and pcDNA3-GSK3β using the lipofectamine method described above. In separate wells, HEK293FT cells were co-transfected with plasmids pRK5-iRFP-tau (alanine mutant) and pcDNA3-GSK3β. Cells were detached from 24-well plate using 0.05 % trypsin-EDTA (Gibco # 25300-054) for 10 min at 37 °C, then pelleted and washed 1x with ice-cold PBSCM (137 mM NaCl, 2.7 mM KCl, 10 mM Na_2_HPO_4_, 1.8 mM KH_2_PO_4_, 1 mM CaCl_2_, and 0.5 mM MgCl_2_, pH 7.4). The same number of cells from the two transfections were mixed and fixed in 4 % paraformaldehyde for 15 min at room temperature in a 1.5 mL microcentrifuge tube with gentle rotating. The fixed cells were permeabilized using PBSCM with 0.1 % Triton X-100 for 15 min at room temperature, then blocked in PBSG (PBSCM with 40 % goat serum, Gibco # 16210-064) for 30 min at room temperature with gentle rotating. For antibody staining, these cells were incubated overnight at 4 °C in dark with gentle rotating with tau antibodies diluted in PBSG (see **Table 1** for dilution). The next day, the cells were incubated with either Alexa Fluor 594 conjugated secondary anti-mouse IgG antibody (1:500, Invitrogen # A11005, RRID: AB_2534073), anti-mouse IgM antibody (1:500, Invitrogen # A21044, RRID: AB_2535713), or anti-rabbit IgG antibody (1:500, Invitrogen # A21442, RRID: AB_2535860) diluted in PBSG for 1 hour at room temperature in dark with continuous rotating. Pelleting and washing 1x with ice-cold PBSCM were required in between each incubation steps. After labeling, the cells were re-suspended in ice-cold PBSCM, and the fluorescence signal from cells was acquired using Becton-Dickinson Fortessa X20 (UConn Center for Open Research Resources and Equipment). The experimenter was not blinded during the experiment.

To measure Φ_λPP_, the HEK293FT cells were co-transfected with plasmids pRK5-EGFP-tau WT and pcDNA3-GSK3β. In separate wells, cells were transfected with pRK5-iRFP-tau WT plasmid only (not co-transfected with pcDNA3-GSK3β), since these cells will be treated with λPP to de-phosphorylate tau. The same number of cells from the two transfections were fixed and permeabilized, separately. After permeabilization, only the cells expressing iRFP-tau WT were treated with 2 U/μL λPP (New England Biolabs # P0753) for 3 hours at 30 °C with gentle rotating. These cells were then mixed with cells expressing EGFP-tau WT. The following blocking, primary and secondary antibody incubation steps follow the same procedure as when measuring Φ_*ala*_, described above. The experimenter was not blinded during the experiment.

### Immunocytochemistry in HEK293FT cells

The HEK293FT cells were seeded in a chamber mounted on #1.0 Borosilicate coverglass (Lab-Tek # 155383), then co-transfected with plasmids pRK5-EGFP-tau WT and pcDNA3-GSK3β as described above. Cells were rinsed 1x with ice-cold PBSCM and fixed in 4 % paraformaldehyde for 15 min at room temperature in the chambers. The fixed cells were permeabilized using PBSCM with 0.1 % Triton X-100 for 15 min at room temperature, blocked in PBSG for 30 min at room temperature, and then incubated with tau antibodies diluted in PBSG overnight at 4 °C in dark. The following tau antibodies and dilutions were used: AT270 (1:200), 1H6L6 (1:250), and AT180 (1:200). The next day, the cells were incubated with either Alexa Fluor 594 conjugated secondary anti-mouse antibody (1:500, Invitrogen # A11005, RRID: AB_2534073) or anti-rabbit antibody (1:500, Invitrogen # A21442, RRID: AB_2535860) diluted in PBSG for 1 hour at room temperature in dark. 300 nM of DAPI was used to stain the nucleus for 1 min at room temperature. 3x washes with ice-cold PBSCM and 5 min incubation on ice in between were required during each incubation steps. The cells were imaged using a Nikon A1R confocal microscope (UConn Center for Open Research Resources and Equipment). The experimenter was not blinded during the experiment.

### Mouse hippocampus dissection

All procedures involving animals were in accordance with the US National Institutes of Health Guide for the Care and Use of Laboratory Animals and approved by the University of Connecticut Institutional Animal Care and Use Committee. Hippocampal dissection was performed according to the methods we have previously described (Cho *et al.* 2019; Klapoetke *et al.* 2014). No sample calculation was performed, but was based on previous studies (Cho *et al.* 2019; Klapoetke *et al.* 2014). Timed pregnant Swiss Webster mice (gestational day 15) were purchased from Taconic. P0 or P1 postnatal mice (randomly selected from litter using simple randomization (Kim & Shin 2014)) were placed on a physical barrier (such as a glove) on top of an ice slurry one at a time to induce hypothermia to reduce or terminate movement and bleeding, and decapitated. Hippocampus was dissected from 6 P0 or P1 postnatal mice in Hank’s balanced salt solution with 10 mM HEPES, 35 mM glucose, 1 mM kynurenic acid, and 10 mM MgCl_2_, pH 7.4. For dissociation, the hippocampal tissue was digested with 50 units of papain (Worthington Biochemical) for 8 min, and halted with ovomucoid trypsin inhibitor (Worthington Biochemical).

### HEK293FT and mouse primary cell lysate preparation and western blotting

The following cell lysates were prepared: non-transfected HEK293FT cells, HEK293FT cells expressing EGFP only, HEK293FT cells expressing EGFP-tau WT, and dissociated mouse hippocampus. Cells were cultured to approximately 80 % confluency on 100 mm polystyrene tissue culture plates and scraped into 1 mL lysis buffer containing 150 mM NaCl, 5 mM EDTA, 50 mM Tris, 1 % Triton X-100, 0.5 % sodium deoxycholate, and 0.1 % sodium dodecyl sulfate, protease inhibitor cocktail (Sigma # P8340), and phosphatase inhibitor cocktails 2&3 (Sigma # 5726 & P0044). Cells were lysed on ice for 15 min and then centrifuged at 13,000 g for 15 min at 4 °C to remove insoluble debris. The concentrations of cell lysate were determined using a Bradford assay kit (ThermoFisher Scientific # 23200) according to the manufacturer’s instructions. The lysates were aliquoted and stored at −20 °C avoiding multiple freeze/thaw cycles.

An equal amount (15 μg) of total cellular protein was separated by SDS-PAGE using 4-12.5 % Bis-Tris gels and transferred to nitrocellulose membrane (GE Healthcare # 10600006). SeeBlue pre-stained protein standard (Invitrogen # LC5925) was used to determine the molecular weight. The nitrocellulose membrane was blocked in TBST (20 mM Tris, 150 mM NaCl, and 0.1 % Tween 20, pH 7.6) with 1 % BSA (bovine serum albumin, ThermoFisher Scientific #149006) for 1 hour at room temperature, and then incubated with the tau antibodies diluted in TBST with 1 % BSA overnight at 4 °C with continuous agitation. The following antibodies and dilutions were used: AT270 (1:1000), 1H6L6 (1:2500), and AT180 (1:1000). The following day, the membrane was washed 3x with TBST and then incubated with either horseradish peroxidase-coupled secondary anti-mouse antibody (1:2000, Invitrogen # 62-6520, RRID: AB_2533947) or anti-rabbit antibody (1:5000, Invitrogen # 31460, RRID: AB_228341) for 1 hour at room temperature with continuous agitation. The peroxidase activity was revealed using an enhanced chemiluminescence detection kit (GE Healthcare, # 16915806). GAPDH antibody (1:2000, Invitrogen # MA5-15738-HRP, RRID: AB_2537659) was used as a loading control. The experimenter was not blinded during the experiment.

### Experimental design

This study was not pre-registered.

## Results

### Phi: A generalizable parameter for measuring the specificity of phospho-tau antibodies

To quantify the specificity of phospho-tau antibodies, we developed a whole-cell based approach using flow cytometry (**Fig. 1**). To probe the site-specificity of phospho-tau antibodies, we generated targeted alanine mutants at defined phosphorylation sites. HEK293FT cells were transiently transfected with either WT human tau (0N4R isoform) fused to the EGFP (EGFP-tau WT), or a point mutant (an alanine mutant at the phosphorylation site of interest) fused to the iRFP (iRFP-tau), in separate wells (**Fig. 1a**). Previous reports indicated that endogenous tau expression is undetected in HEK293 cells (Kfoury *et al.* 2012; Liu *et al.* 2013). In addition, when transfected with human tau, endogenous kinases in HEK293FT cells phosphorylated human tau (Camero *et al.* 2014; Houck *et al.* 2016). Leveraging these facts, we developed a ratiometric assay to quantify the specificity of phospho-tau antibodies (**Fig. 1a**). The EGFP-positive cells (expressing WT tau) and iRFP-positive cells (expressing a tau point mutant incapable of phosphorylation at a defined site) were mixed in same numbers and incubated with phospho-tau antibodies (**Fig. 1a**). The EGFP (green channel) and iRFP (far-red channel) positive cells, and their antibody binding (red channel) were clearly distinguished using flow cytometry (**Fig. 1b**, left panel). This assay quantifies the specific binding of the antibody to a defined phosphorylation site. In this assay, the phospho-tau antibody AT180, targeting phospho-threonine 231 (pThr231) (Goedert *et al.* 1994; Gotz *et al.* 2001; Nakamura *et al.* 2012), showed almost exclusive labeling to the cells expressing EGFP-tau WT, and no detectable binding to iRFP-tau T231A (**Fig. 1b**), indicating that AT180 binding requires pThr231. In case an antibody shows non-specific binding to the alanine mutant at the target modification site, the antibody may be recognizing a phosphorylated residue at an off-target site. In addition, the absence of antibody binding to the alanine mutant does not exclude the possibility that binding to the non-modified tau is inhibited by the alanine mutation. In other words, the antibody may possess non-specific binding to unmodified tau, but may not bind to the alanine point mutant. To assess these possibilities, we transfected HEK293FT cells with either EGFP-tau WT or iRFP-tau WT separately and treated only the iRFP-tau WT expressing cells with λPP (**Fig. 1a**). As in the assay using the T231A mutant, AT180 showed exclusive binding to EGFP-tau WT cells (**Fig. 1b**, right panel). This result provides further support that the AT180 antibody does not recognize the WT tau epitope without phosphorylation of Thr231.

We tested the assay (**Fig. 1a)** for its ability to discern phospho-specificity using 3 previously characterized anti-tau antibodies. Tau-5 is a pan-tau antibody that shows phosphorylation-independent binding (LoPresti *et al.* 1995; Carmel *et al.* 1996), 3.24 is a high specificity scFv to pThr231 tau (Li *et al.* 2018), and ab131354 targets pSer262 tau but have shown cross-reactivity to non-phosphorylated tau (Ercan *et al.* 2017). Tau-5 indeed showed binding to both EGFP-tau WT and iRFP-tau WT treated with λPP (**Fig. 2a**). 3.24 showed binding to EGFP-tau WT, but not to iRFP-tau T231A or iRFP-tau WT treated with λPP (**Fig. 2b**), confirming its high specificity. ab131354 showed cross-reactivity to both iRFP-tau S262A and iRFP-tau WT treated with λPP (**Fig. 2c**), indicating that it is not specific to pSer262 tau.

**Figure 2.**
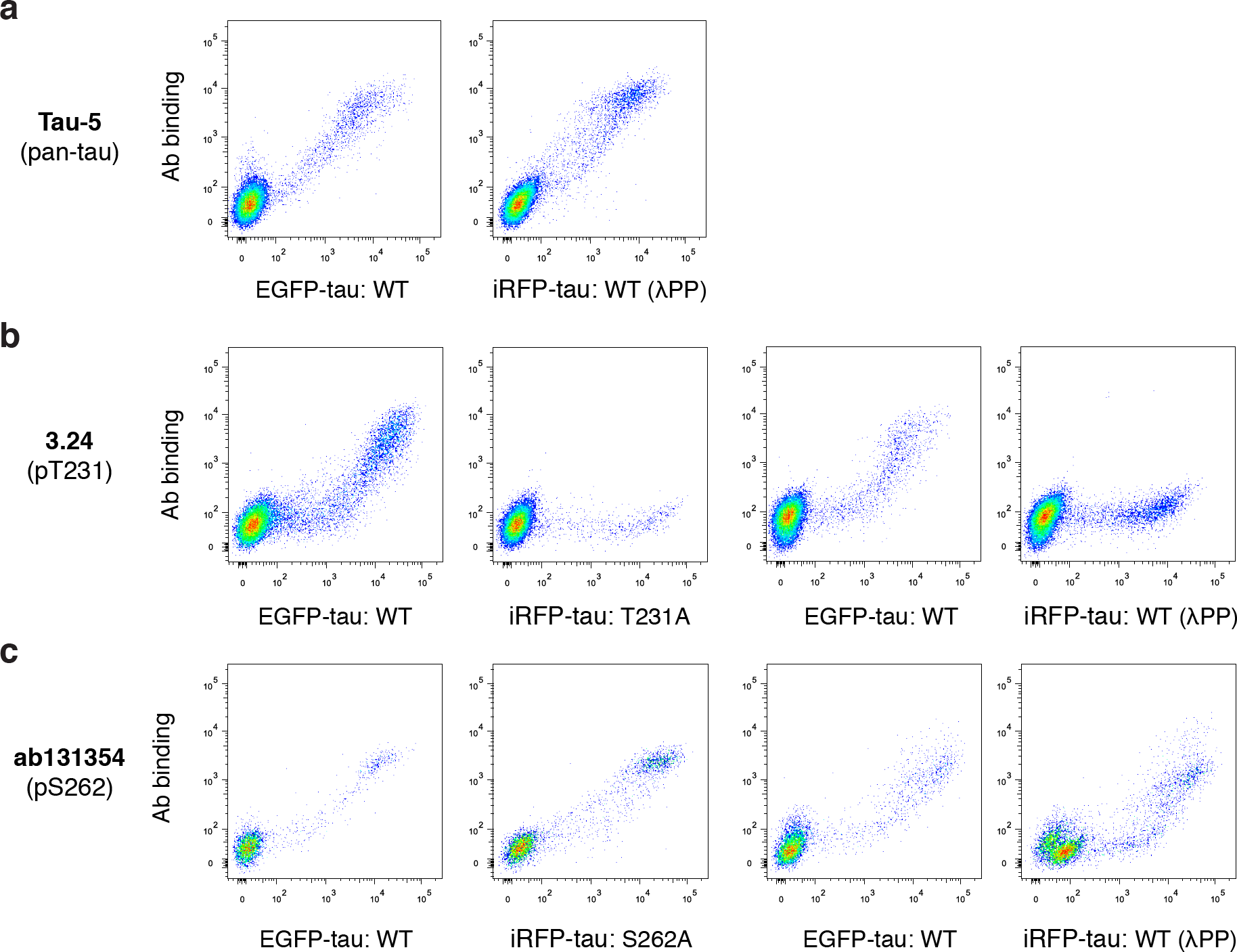
Validation of the assay using well characterized anti-tau antibodies. **a**, Flow cytometry plots for the pan-tau antibody Tau-5, which shows binding to both EGFP-tau WT and iRFP-tau WT treated with λPP. **b-c**, Flow cytometry plots for the high specificity single-chain variable fragment 3.24 (**b)** and ab131354 (**c)**.

Based on this assay, we introduce a parameter termed Φ (see Materials and Methods section for definition) that quantifies the specificity of phospho-site targeting antibodies. Φ is determined by the degree of cross-reactivity towards cells expressing the target antigen with an alanine point mutation at the target phospho-site, or cells expressing the target antigen but were treated with λPP for de-phosphorylation. Φ ranges from 0 to 1, where 1 indicates perfect specificity (no binding to the alanine mutant or λPP treated cells), and lower Φ indicates more false positive detection of nonphospho-tau or phospho-tau at off-target sites. 2 Φ values are derived from the definition: Φ_ala_ is determined from cross-reactivity to tau with a point alanine mutation at the target site, while Φ_λPP_ is determined by cross-reactivity to de-phosphorylated tau. The phospho-site specificity is determined by evaluating both Φ_ala_ and Φ_λPP_ values. The specificity Φ_ala_ and Φ_λPP_ of AT180 were 0.99 ± 0.01 and 0.98 ± 0.01, respectively (**Table 1**, mean ± standard deviation from 2 independent experiments, all data points shown in **Supp. Fig. 1**), indicating nearly perfect phospho-site specificity towards pThr231 tau. The specificity Φ_ala_ and Φ_λPP_ of ab131354 were 0.32 ± 0.02 and 0.37 ± 0.01, respectively, reflecting its poor specificity (**Fig. 2**). Φ can be used as a general standard to measure and compare the specificity of post-translational modification (PTM)-site targeting antibodies.

### Co-expression of glycogen synthase kinase-3β enhances tau phosphorylation

The assay developed here relies on intracellular phosphorylation of tau due to endogenous kinase activity in HEK293FT cells. Previous studies have found that tau expressed in HEK293 cells were phosphorylated at various sites, including S214, S262, S396/S404 (Ren *et al.* 2007). A major advantage of our assay is the potential to co-express kinases to induce tau phosphorylation. Since many of the residues that are phosphorylated on tau are serine or threonine preceding a proline, we co-expressed glycogen synthase kinase (GSK)-3β, a proline-directed kinase, in HEK293FT cells to enhance tau phosphorylation. GSK-3β phosphorylates tau at a wide range of sites found in Alzheimer’s disease patients (Mandelkow *et al.* 1992). Co-transfected cells showed a similar level of EGFP-tau WT expression (**Fig. 3a, b**), but enhanced binding to the phospho-tau antibody AT8, which targets pSer202 (Pei *et al.* 1999; Sereno *et al.* 2009; Hernandez *et al.* 2003) (**Fig. 3b**), indicating improved phosphorylation. Therefore, to enhance the detection of phospho-tau antibody binding, we co-transfected tau constructs with GSK-3β in subsequent experiments.

**Figure 3.**
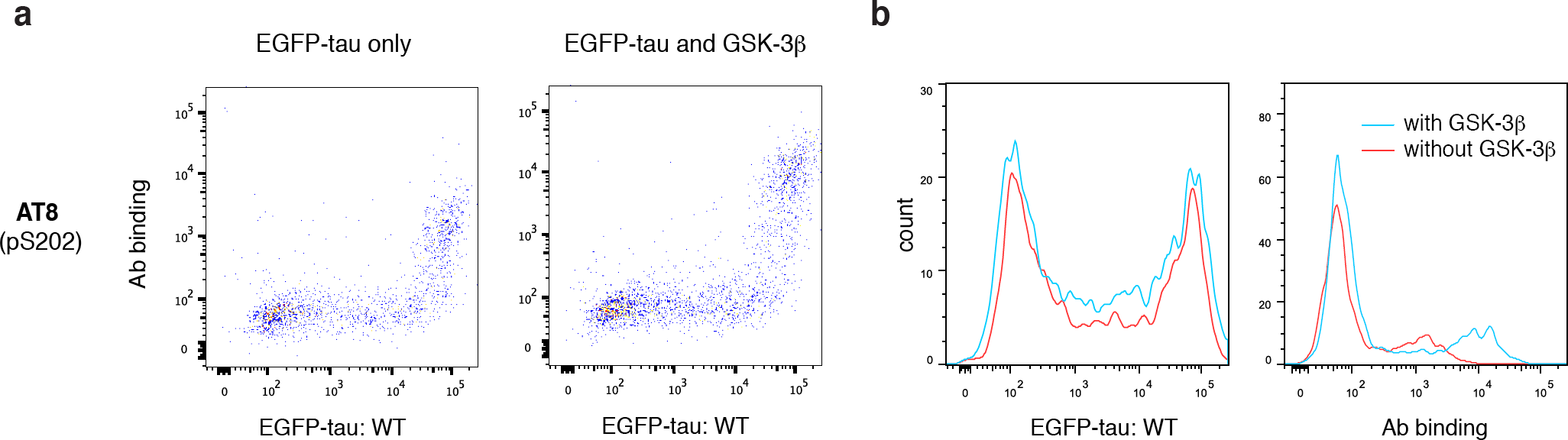
Co-expression of GSK-3β in HEK293FT cells enhances phosphorylation of tau. **a**, Representative flow cytometry plots for AT8 showing tau expression (EGFP-tau) and AT8 binding cells transfected with tau-only (left) or cells co-transfected with tau and GSK-3β (right). **b**, Histograms for tau expression (left) and AT8 binding (right) in cells transfected with tau-only (red) or cells co-transfected with tau and GSK-3β (blue).

### Quantifying the phospho-site specificity of commonly used phospho-tau antibodies

Using our assay, we quantified the specificity Φ of 7 widely used phospho-tau antibodies (AT270, AT8, AT100, AT180, PHF-6, TG-3, and PHF-1) as well as 2 additional phospho-tau antibodies (1H6L6 and pS404). Collectively these antibodies have been raised against 6 distinct phosphorylation sites (see **Table 1** for more information) of high relevance in a wide range of neurodegeneration. We applied the same approach described in **Fig. 1** to quantify both Φ_ala_ and Φ_λPP_. In **Fig. 4**, data in the left two columns show antibody labeling from a mixture of cells transfected with either EGFP-tau WT or iRFP-tau with the corresponding alanine point mutant. The right two columns show antibody labeling from cells transfected with either EGFP-tau WT or iRFP-tau WT treated with λPP. Clear binding to EGFP-tau WT was detected in all antibodies except AT100 (**Fig. 4**, leftmost column). Most antibodies showed exclusive binding to EGFP-tau WT, and no binding to the alanine point mutant at their corresponding target site or the iRFP-tau WT treated with λPP (**Fig. 4**, second and last columns from left). From two independent experiments, we obtained highly reproducible measurement of Φ_ala_ and Φ_λPP_ from these antibodies (**Table 1** and **Supp. Fig. 1**). 5 out of 7 antibodies (AT8, AT180, PHF-6, TG-3, and PHF-1) had Φ values near 1, indicating these antibodies have nearly perfect specificity. For AT270, Φ_ala_ = 0.80 ± 0.06 (**Table 1**), due to the elevated level of binding signal in cells expressing the T181A mutant (**Fig. 4**). However, Φ_λPP_= 0.99 ± 0.01 (**Table 1**), indicating that the cross-reactivity was not detected in dephosphorylated cells. Taken Φ_*ala*_ and Φ_λPP_ together, the non-specific signal detected from AT270 was phosphorylation-sensitive (see below for further characterization of non-specific binding in AT270). For AT100, the absence of binding to EGFP-tau WT cells suggests that the phospho-site (pThr212/pSer214) was not phosphorylated in HEK293FT cells, even with co-transfection of GSK-3β. Therefore, we were unable to measure the Φ values for AT100. Taken together, these results validate that the widely used phospho-tau antibodies AT8, AT180, PHF-6, TG-3, and PHF1 binds specifically to tau phosphorylated at the target site, but not to non-phosphorylated tau or phosphorylated tau at off-target sites.

**Figure 4.**
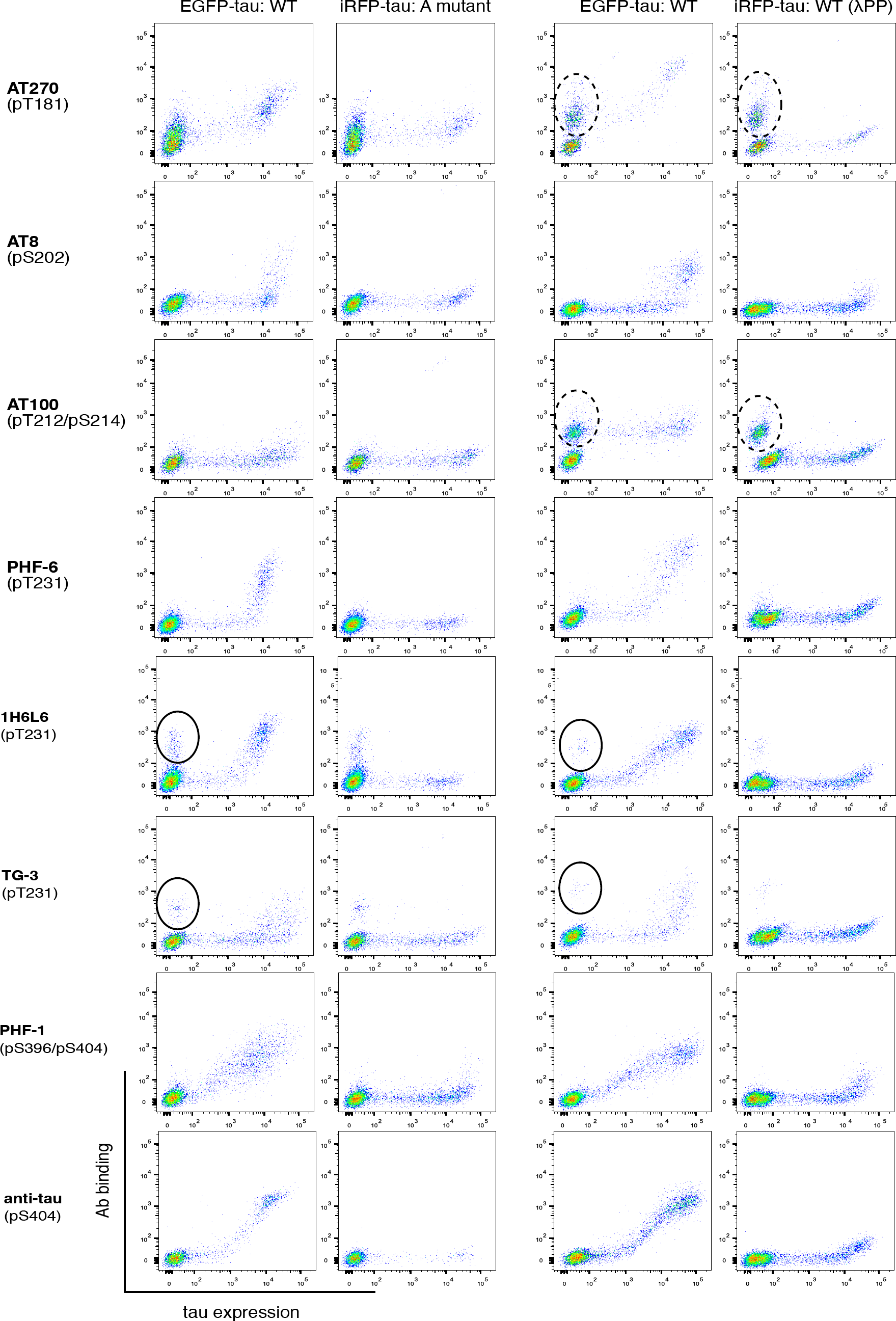
Specificity measurements for commonly used phospho-tau antibodies using the flow cytometry assay described in Fig. 1. Dotted and closed circles indicate observed non-specific binding to cells not expressing tau.

### Non-specific binding to cells in phospho-tau antibodies

For the antibody AT270, which showed lower Φ_ala_ but high Φ _λPP_, we observed a reproducible non-specific binding to cells not expressing tau (dotted circles in **Fig. 4**). Similar non-specific binding was detected with AT100, even though AT100 did not bind to EGFP-tau WT. Although not as prominent, a small fraction and weak level of non-specific binding to tau-negative cells was detected in antibodies 1H6L6 and TG-3 (closed circles in **Fig. 4**). For other antibodies, background binding to non-transfected cells was not detected. The binding to cells not expressing tau indicates that these antibodies exhibit non-specific binding to HEK293FT cells. We conducted experiments to rule out the possibility that the non-specific binding is originating from the secondary reagents used for detection. Control antibodies having matched isotype (mouse IgG1, mouse IgM, or rabbit IgG) followed by the secondary antibodies (see Materials and Methods section) were used to label a mixture of cells transfected with either EGFP-tau WT, or cells transfected with iRFP-tau WT treated with λPP (**Supp. Fig. 2**). No non-specific binding was detected in the secondary reagents, indicating that the non-specific binding observed was not due to secondary reagents. For AT270 and AT100, the population with non-specific binding separated clearly when cells were treated with λPP (**Fig. 4**, right two columns). Based on this result, we hypothesized that the non-specific bindings of AT270 and AT100 are due to cross-reactivity to other phosphorylated proteins in HEK293FT cells. In this scenario, cells treated with λPP would show decreased level of antibody binding, while untreated cells would have generally elevated antibody binding. To test this, we labeled a mixture of cells transfected with either EGFP (not EGFP-tau), or cells transfected with iRFP (not iRFP-tau) treated with λPP. In this experiment, binding was detected only in GFP-positive cells (**Fig. 5a**). Since specific binding to phospho-tau in AT100 was not detected, we further analyzed the non-specific binding of AT270. We also analyzed the non-specific binding of 1H6L6, which showed a weak level of non-specific binding (closed circles in **Fig. 4**). TG-3 was not included in this analysis since the fraction of tau-negative cells showing non-specific binding was even smaller. To further assess the non-specific binding found in AT270 and 1H6L6, we conducted immunocytochemistry experiments. Antibodies AT270 and 1H6L6 showed binding in cells not expressing EGFP-tau, while the highly specific antibody AT180 showed staining only to EGFP-tau positive cells (**Fig. 5b**). To assess the broadness of proteins that cause the non-specific binding, we conducted western blotting experiments. We probed cell lysates from non-transfected, EGFP-transfected, EGFP-tau transfected cells as well as dissociated mouse hippocampus. In these experiments, antibodies AT270 and 1H6L6 showed bands in addition to a band corresponding to EGFP-tau (**Fig. 5c**). In contrast, a single band corresponding to EGFP-tau was observed in AT180 (**Fig. 5c**). For AT270, off-target bands found in HEK293FT cell lysates were also detected in WT mouse hippocampal cell lysate. Therefore, this non-specific binding was independent of species, leading to a non-specific band interfering with the mouse tau signal. However, for 1H6L6, the non-specific binding was not clearly detected in mouse hippocampal lysate, suggesting that the non-specific binding may be species or cell-type dependent. These results demonstrate that by using a whole-cell based approach, we have identified a source of non-specific binding in antibodies AT270 and 1H6L6 not observable when using peptides for specificity characterization.

**Figure 5.**
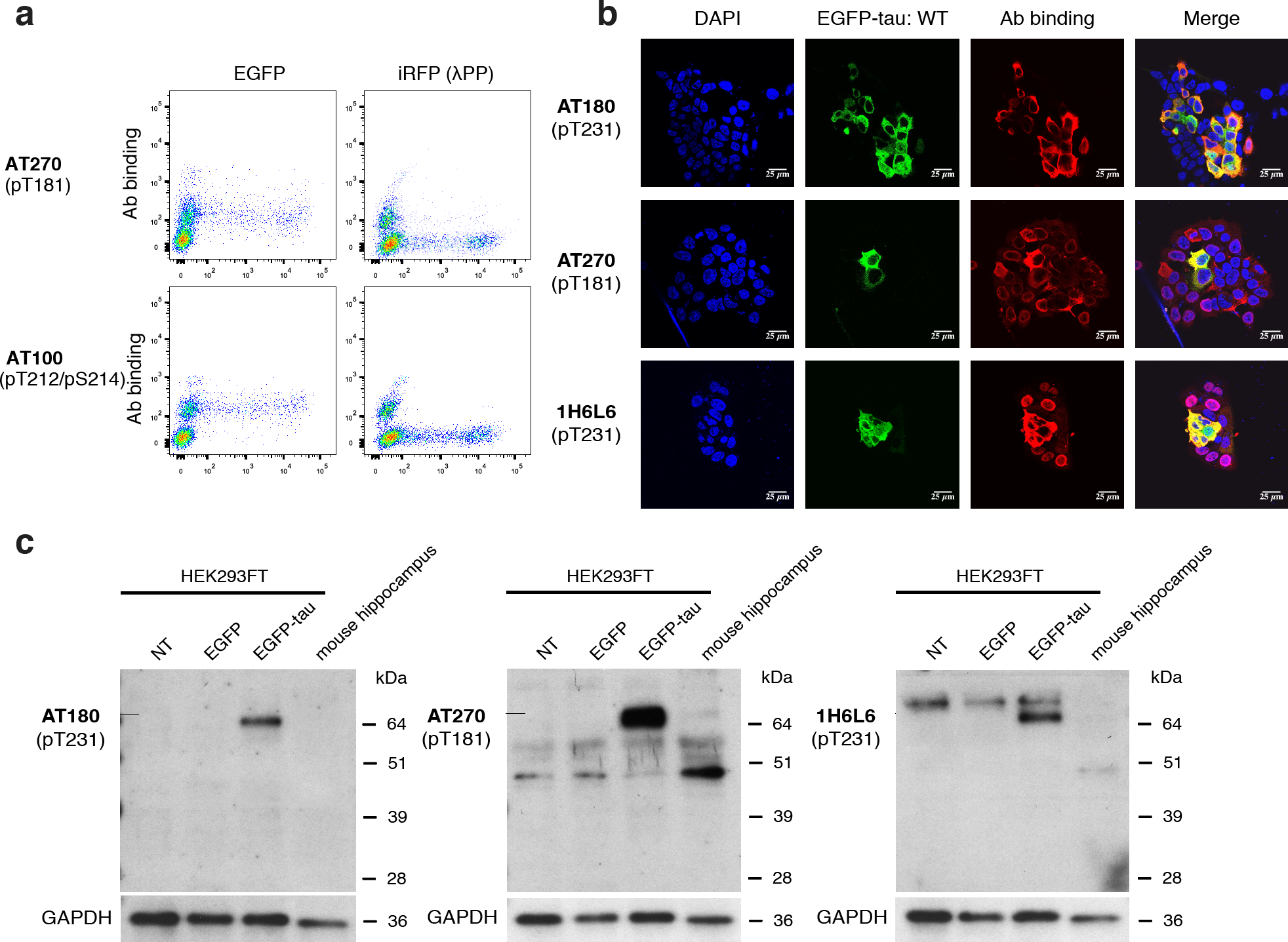
Non-specific binding to HEK293FT cells in phospho-tau antibodies. **a**, Flow cytometry plots for AT270 and AT100 to detect non-specific binding to normal HEK293FT cells (EGFP-transfected) vs. λPP-treated cells (iRFP-transfected). **b**, Confocal images of AT180, AT270, and 1H6L6 antibodies binding to HEK293FT cells transfected with EGFP-tau WT. Images show nuclei stained using DAPI (blue), tau expression (green), antibody staining (red), and merge. Scale bars, 25 μm. **c**, Analysis of tau signal and non-specific signal in AT270 and 1H6L6 by western blotting. The proteins were extracted from: lane 1, non-transfected HEK293FT cells; lane 2, HEK293FT cells transfected with EGFP; lane 3, HEK293FT cells co-transfected with EGFP-Tau/GSK-3β; lane 4, mouse hippocampus. Proteins were identified with the following antibodies: AT180, AT270, and 1H6L6. GAPDH was used as a loading control.

## Discussion

Site-specific phosphorylation of tau is critical in understanding the molecular biology of tau and is a well-defined target in neurodegeneration. The specificity of phospho-tau antibodies is critical for reproducible and sensitive detection (Li *et al.* 2018; Lothrop *et al.* 2013). Phospho-tau antibodies should interact only with the target epitope sequence with site-specific phosphorylation, and not with nonphospho-tau or phospho-tau at off-target sites. In particular, considering the fact that phospho-tau antibodies are being developed for diagnostic tests and clinical trials as therapeutics, a standard measure of specificity is urgently needed (Chai *et al.* 2011; Boutajangout *et al.* 2011; Kayed 2010; Asuni *et al.* 2007; Boutajangout *et al.* 2010; Lewis *et al.* 2000) In this study, we developed a whole-cell based assay that enables robust measurement of the specificity of phospho-tau antibodies, and introduced the specificity parameter termed Φ. Φ is determined by measuring the fraction of binding signal from cross-reactivity to nonphospho-tau or off-target phosphorylation sites. Φ ranges from 0 to 1, where 1 indicates perfect specificity and lower Φ indicates more false positive detection of either nonphospho-tau sequences or the phospho-tau at the off-target site. We measured two Φ values (Φ_ala_ and Φ_λPP_) to comprehensively capture the specificity. Φ_ala_ indicates the phospho-site specificity of the antibody by measuring the fraction of non-specific binding to an alanine point mutant at the target phospho-site. Φ_λPP_ indicates the phospho-sensitivity of the antibody by measuring non-specific binding to de-phosphorylated tau. This approach can be easily adapted to other phosphorylated proteins, and potentially to other PTMs such as methylation and acetylation, allowing specificity quantification and comparison of any PTM-site specific antibody. In addition, the Φ values can be combined to a single parameter, to yield a single specificity parameter. For example,

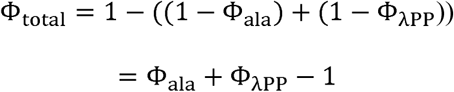

In our calculation of the Φ values, we did not include the fluorescence signal detected from cells not expressing tau. Therefore, the small non-specific binding to cells we observed for antibody 1H6L6 (and to an even smaller degree in TG-3) is not reflected in their Φ values. As for AT270, the Φ value captured the non-specific binding to cells, since the non-specific binding was detected in the cells expressing the alanine mutant but was lost after de-phosphorylation with λPP. We attempted to define another Φ parameter to quantify the non-specific binding to cells but the resulting values were inconsistent due to variations in the level of non-specific binding signal and fraction of cells showing binding.

Using our assay, we measured the Φ values of 10 phospho-tau antibodies, 7 of which have been used in hundreds of previous studies. Out of the 7, we quantified the specificity of 6 antibodies (AT270, AT8, AT180, PHF-6, TG-3, and PHF-1). We were unable to determine Φ values for AT100, due to lack of binding to tau expressed in HEK293FT cells. Endogenous kinases in HEK293FT cells phosphorylate human tau (Camero *et al.* 2014; Houck *et al.* 2016). We showed that co-transfection of tau with GSK-3β led to increased phosphorylation of tau, but not for the AT100 target site (Thr212/Ser214). Previous studies have indicated that the AT100 epitope is formed only when a specific order of phosphorylation occurs (Zheng-Fischhofer *et al.* 1998). It has also been shown that Thr212 and Ser214 are sequentially phosphorylated by GSK-3β and protein kinase A (PKA), respectively (Zheng-Fischhofer *et al.* 1998). Considering that PKA is commonly expressed in cells, tau expressed in HEK293FT cells may only be phosphorylated at Ser214, leading to lack of AT100 binding. Tau phosphorylation may be enhanced by using mammalian brain extracts, which contain kinases that phosphorylate tau *in vitro* (Gustke *et al.* 1992; Lichtenberg-Kraag *et al.* 1992; Zheng-Fischhofer *et al.* 1998).

Out of the 6 phospho-tau antibodies (AT270, AT8, AT180, PHF-6, TG-3, and PHF-1), all but AT270 had Φ values close to 1, indicating near-perfect specificity. For AT270, a weak, but reproducible binding was detected to tau-negative HEK293FT cells (**Fig. 4**). This non-specific binding was consistently detected in immunocytochemistry experiments, leading to distinct perinuclear non-specific binding (**Fig. 5b**). Western blotting with AT270 revealed additional bands consistently present in non-transfected HEK293FT cell lysate as well as primary WT mouse hippocampal lysate (**Fig. 5c**). A previous study found multiple bands when probing WT mouse brain lysate (Petry *et al.* 2014). The same study also found non-specific binding in antibodies AT8, AT180, and TG-3, using cell lysate prepared from tau knockout mice. However, this non-specific binding was not due to the antibodies themselves, and was eliminated by switching secondary reagents (Petry *et al.* 2014). We found that the two Φ values for AT270 was inconsistent (Φ_ala_ = 0.80 ± 0.06 vs. Φ_λPP_= 0.99 ± 0.01). This indicated that the source of cross-reactivity was lost upon de-phosphorylation with λPP, suggesting that AT270 may bind to another phosphorylated protein. Indeed, the non-specific binding was only detected from the tau-negative cells expressing EGFP, but not from the tau-negative cells expressing iRFP with de-phosphorylation treatment (**Fig. 5a**).

A major advantage of our specificity assay is the use of flow cytometry, which allows sensitive, quantitative detection of antibody binding. We enabled the distinction of non-specific binding within a single sample by tagging potential cross-reactive proteins with a fluorescent protein easily distinguishable from the WT target protein. These features led to the reproducible measurement of Φ values for a wide range of phospho-tau antibodies. Previously, peptide arrays have been used to validate the specificity of PTM-specific antibodies, such as those targeting histone modifications (Egelhofer *et al.* 2011; Bock *et al.* 2011; Fuchs *et al.* 2011). Synthetic peptides and peptide arrays were also used for specificity analysis of phospho-tau antibodies (Ercan *et al.* 2017). The limitation of this approach is the high cost of synthesizing peptides in order to comprehensively cover tau phosphorylation sites. Moreover, short peptides do not have strong binding to conformation-sensitive phospho-tau antibodies, such as TG-3 (Gotz *et al.* 2001). Our approach allows assessment of binding to full-length tau with defined sequence variations, and can be co-expressed with kinases to induce tau phosphorylation. Using our approach, the specificity of the conformation-sensitive pT231 tau antibody TG-3 was also successfully determined (**Fig. 4** and **Table 1**). Libraries of tau mutations may be generated, and deep sequencing approaches can be used for *de novo* identification of anti-tau antibodies. Our approach also revealed non-specific binding of antibodies to irrelevant cellular proteins. We found that the source of non-specific binding for AT270 is conserved across HEK293FT cells and mouse primary hippocampal cells, but was not conserved for 1H6L6, indicating cell-type and species sensitivity of non-specific binding. Further analyses using immunoprecipitation and mass spectrometry may allow the identification of the cross-reactive proteins.

Since the cross-reactivity of antibodies would be dependent on the affinity to the off-target sites or molecules, it is possible that the Φ values are dependent on antibody concentration. In particular, the Φ values may decrease upon increasing antibody concentration. In our experiments, we used antibody concentrations typically used for cell labeling experiments (2 μg/mL, **Table 1**). Since we observed high antibody binding signal in these experiments for all antibodies tested except AT100 (which showed non-specific binding to cells), we did not attempt to increase the antibody concentration. Also, all antibodies that showed binding had near-perfect specificity, except AT270 and ab131354. Lowering the antibody concentration for these antibodies resulted in a significant decrease in antibody binding, and did not result in improvement of Φ values (data not shown). Even for the high specificity antibodies, users should be cautious when using them at higher concentrations than what is listed in **Table 1**, since that may result in increased non-specific binding.

In summary, our study provided a whole-cell based approach to validate the specificity of phospho-tau antibodies. Using this assay, we determined the parameter Φ for phospho-tau antibodies, that enable quantitative comparison of specificity. We anticipate that this parameter will be widely adopted in guiding users of the specificity of phospho-tau antibodies, and potentially PTM-specific antibodies in general.

## List of abbreviations

AD: Alzheimer’s disease
HEK: Human embryonic kidney
phospho-tau: phosphorylated tau
PHF: paired helical filament
EGFP: enhanced green fluorescent protein
iRFP: infrared fluorescent protein
λPP: lambda protein phosphatase
WT: wild-type
GSK-3β: glycogen synthase kinase-3β
PTM: post-translational modification
scFv: single-chain antibody variable fragment
pThr231: phospho-threonine 231
RRID: Research Resource Identifier

## Acknowledgements

This work was funded by the NIH grant NS099903 and NSF grant 1706743. D.L. was also supported by the UConn Institute for Systems Genomics Munson award, and the UConn-Alexion student dissertation improvement award.

## Supplementary Figures

**Supplementary Figure 1.** Measurement of Φ_ala_ and Φ_λPP_ of phospho-tau antibodies. Error bars indicate standard deviation from two independent cell culture preparations. Red, Φ_λPP_. Blue, Φ_ala_.

**Supplementary Figure 2.** Flow cytometry plots showing binding of isotype-matching control antibodies under the specificity quantification assay conditions.

## References

Asuni, A. A., Boutajangout, A., Quartermain, D. and Sigurdsson, E. M. (2007) Immunotherapy targeting pathological tau conformers in a tangle mouse model reduces brain pathology with associated functional improvements. J Neurosci 27, 9115–9129.

Bock, I., Dhayalan, A., Kudithipudi, S., Brandt, O., Rathert, P. and Jeltsch, A. (2011) Detailed specificity analysis of antibodies binding to modified histone tails with peptide arrays. Epigenetics 6, 256–263.

Boutajangout, A., Ingadottir, J., Davies, P. and Sigurdsson, E. M. (2011) Passive immunization targeting pathological phospho-tau protein in a mouse model reduces functional decline and clears tau aggregates from the brain. Journal of neurochemistry 118, 658–667.

Boutajangout, A., Quartermain, D. and Sigurdsson, E. M. (2010) Immunotherapy targeting pathological tau prevents cognitive decline in a new tangle mouse model. J Neurosci 30, 16559–16566.

Caccamo, A., Majumder, S., Richardson, A., Strong, R. and Oddo, S. (2010) Molecular interplay between mammalian target of rapamycin (mTOR), amyloid-beta, and Tau: effects on cognitive impairments. J Biol Chem 285, 13107–13120.

Camero, S., Benitez, M. J., Cuadros, R., Hernandez, F., Avila, J. and Jimenez, J. S. (2014) Thermodynamics of the interaction between Alzheimer’s disease related tau protein and DNA. PLoS One 9, e104690.

Carmel, G., Mager, E. M., Binder, L. I. and Kuret, J. (1996) The structural basis of monoclonal antibody Alz50’s selectivity for Alzheimer’s disease pathology. The Journal of biological chemistry 271, 32789–32795.

Chai, X., Wu, S., Murray, T. K. et al. (2011) Passive immunization with anti-Tau antibodies in two transgenic models: reduction of Tau pathology and delay of disease progression. J Biol Chem 286, 34457–34467.

Cho, Y. K., Park, D., Yang, A., Chen, F., Chuong, A. S., Klapoetke, N. C. and Boyden, E. S. (2019) Multidimensional screening yields channelrhodopsin variants having improved photocurrent and order-of-magnitude reductions in calcium and proton currents. The Journal of biological chemistry 294, 3806–3821.

Dengler-Crish, C. M., Smith, M. A. and Wilson, G. N. (2017) Early Evidence of Low Bone Density and Decreased Serotonergic Synthesis in the Dorsal Raphe of a Tauopathy Model of Alzheimer’s Disease. J Alzheimers Dis 55, 1605–1619.

Dickson, D. W., Crystal, H. A., Bevona, C., Honer, W., Vincent, I. and Davies, P. (1995) Correlations of synaptic and pathological markers with cognition of the elderly. Neurobiol Aging 16, 285–298; discussion 298-304.

Egelhofer, T. A., Minoda, A., Klugman, S. et al. (2011) An assessment of histone-modification antibody quality. Nature structural & molecular biology 18, 91–93.

Engler, C., Kandzia, R. and Marillonnet, S. (2008) A one pot, one step, precision cloning method with high throughput capability. PloS one 3, e3647.

Ercan, E., Eid, S., Weber, C. et al. (2017) A validated antibody panel for the characterization of tau post-translational modifications. Mol Neurodegener 12, 87.

Fuchs, S. M., Krajewski, K., Baker, R. W., Miller, V. L. and Strahl, B. D. (2011) Influence of combinatorial histone modifications on antibody and effector protein recognition. Curr Biol 21, 53–58.

Goedert, M., Jakes, R., Crowther, R. A., Cohen, P., Vanmechelen, E., Vandermeeren, M. and Cras, P. (1994) Epitope mapping of monoclonal antibodies to the paired helical filaments of Alzheimer’s disease: identification of phosphorylation sites in tau protein. Biochem J 301 (Pt 3), 871–877.

Goedert, M., Jakes, R. and Vanmechelen, E. (1995) Monoclonal antibody AT8 recognises tau protein phosphorylated at both serine 202 and threonine 205. Neurosci Lett 189, 167–169.

Gotz, J., Chen, F., Barmettler, R. and Nitsch, R. M. (2001) Tau filament formation in transgenic mice expressing P301L tau. J Biol Chem 276, 529–534.

Gotz, J., Probst, A., Spillantini, M. G., Schafer, T., Jakes, R., Burki, K. and Goedert, M. (1995) Somatodendritic localization and hyperphosphorylation of tau protein in transgenic mice expressing the longest human brain tau isoform. EMBO J 14, 1304–1313.

Greenberg, S. G., Davies, P., Schein, J. D. and Binder, L. I. (1992) Hydrofluoric acid-treated tau PHF proteins display the same biochemical properties as normal tau. J Biol Chem 267, 564–569.

Gustke, N., Steiner, B., Mandelkow, E. M., Biernat, J., Meyer, H. E., Goedert, M. and Mandelkow, E. (1992) The Alzheimer-like phosphorylation of tau protein reduces microtubule binding and involves Ser-Pro and Thr-Pro motifs. FEBS Lett 307, 199–205.

Hall, S. S. and Daugherty, P. S. (2009) Quantitative specificity-based display library screening identifies determinants of antibody-epitope binding specificity. Protein science: a publication of the Protein Society 18, 1926–1934.

Hernandez, F., Lucas, J. J., Cuadros, R. and Avila, J. (2003) GSK-3 dependent phosphoepitopes recognized by PHF-1 and AT-8 antibodies are present in different tau isoforms. Neurobiol Aging 24, 1087–1094.

Hoffmann, R., Lee, V. M., Leight, S., Varga, I. and Otvos, L., Jr. (1997) Unique Alzheimer’s disease paired helical filament specific epitopes involve double phosphorylation at specific sites. Biochemistry 36, 8114–8124.

Houck, A. L., Hernandez, F. and Avila, J. (2016) A Simple Model to Study Tau Pathology. J Exp Neurosci 10, 31–38.

Jackson, G. R., Wiedau-Pazos, M., Sang, T. K., Wagle, N., Brown, C. A., Massachi, S. and Geschwind, D. H. (2002) Human wild-type tau interacts with wingless pathway components and produces neurofibrillary pathology in Drosophila. Neuron 34, 509–519.

Janes, K. A. (2015) An analysis of critical factors for quantitative immunoblotting. Sci Signal 8, rs2.

Jicha, G. A., Lane, E., Vincent, I., Otvos, L., Jr., Hoffmann, R. and Davies, P. (1997) A conformation- and phosphorylation-dependent antibody recognizing the paired helical filaments of Alzheimer’s disease. Journal of neurochemistry 69, 2087–2095.

Kayed, R. (2010) Anti-tau oligomers passive vaccination for the treatment of Alzheimer disease. Hum Vaccin 6, 931–935.

Kfoury, N., Holmes, B. B., Jiang, H., Holtzman, D. M. and Diamond, M. I. (2012) Trans-cellular propagation of Tau aggregation by fibrillar species. The Journal of biological chemistry 287, 19440–19451.

Kim, J. and Shin, W. (2014) How to do random allocation (randomization). Clin Orthop Surg 6, 103–109.

Klapoetke, N. C., Murata, Y., Kim, S. S. et al. (2014) Independent optical excitation of distinct neural populations. Nature methods 11, 338–346.

Lewis, J., McGowan, E., Rockwood, J. et al. (2000) Neurofibrillary tangles, amyotrophy and progressive motor disturbance in mice expressing mutant (P301L) tau protein. Nat Genet 25, 402–405.

Li, D., Wang, L., Maziuk, B. F., Yao, X., Wolozin, B. and Cho, Y. K. (2018) Directed evolution of a picomolar-affinity, high-specificity antibody targeting phosphorylated tau. The Journal of biological chemistry 293, 12081–12094.

Lichtenberg-Kraag, B., Mandelkow, E. M., Biernat, J., Steiner, B., Schroter, C., Gustke, N., Meyer, H. E. and Mandelkow, E. (1992) Phosphorylation-dependent epitopes of neurofilament antibodies on tau protein and relationship with Alzheimer tau. Proc Natl Acad Sci U S A 89, 5384–5388.

Liu, Y., Su, Y., Wang, J., Sun, S., Wang, T., Qiao, X., Run, X., Li, H. and Liang, Z. (2013) Rapamycin decreases tau phosphorylation at Ser214 through regulation of cAMP-dependent kinase. Neurochemistry international 62, 458–467.

LoPresti, P., Szuchet, S., Papasozomenos, S. C., Zinkowski, R. P. and Binder, L. I. (1995) Functional implications for the microtubule-associated protein tau: localization in oligodendrocytes. Proc Natl Acad Sci U S A 92, 10369–10373.

Lothrop, A. P., Torres, M. P. and Fuchs, S. M. (2013) Deciphering post-translational modification codes. FEBS Lett 587, 1247–1257.

Mandelkow, E. M., Drewes, G., Biernat, J., Gustke, N., Van Lint, J., Vandenheede, J. R. and Mandelkow, E. (1992) Glycogen synthase kinase-3 and the Alzheimer-like state of microtubule-associated protein tau. FEBS Lett 314, 315–321.

Mercken, M., Vandermeeren, M., Lubke, U., Six, J., Boons, J., Van de Voorde, A., Martin, J. J. and Gheuens, J. (1992) Monoclonal antibodies with selective specificity for Alzheimer Tau are directed against phosphatase-sensitive epitopes. Acta Neuropathol 84, 265–272.

Nakamura, K., Greenwood, A., Binder, L., Bigio, E. H., Denial, S., Nicholson, L., Zhou, X. Z. and Lu, K. P. (2012) Proline isomer-specific antibodies reveal the early pathogenic tau conformation in Alzheimer’s disease. Cell 149, 232–244.

Otvos, L., Jr., Feiner, L., Lang, E., Szendrei, G. I., Goedert, M. and Lee, V. M. (1994) Monoclonal antibody PHF-1 recognizes tau protein phosphorylated at serine residues 396 and 404. Journal of neuroscience research 39, 669–673.

Pavoor, T. V., Cho, Y. K. and Shusta, E. V. (2009) Development of GFP-based biosensors possessing the binding properties of antibodies. Proceedings of the National Academy of Sciences of the United States of America 106, 11895–11900.

Pei, J. J., Braak, E., Braak, H., Grundke-Iqbal, I., Iqbal, K., Winblad, B. and Cowburn, R. F. (1999) Distribution of active glycogen synthase kinase 3beta (GSK-3beta) in brains staged for Alzheimer disease neurofibrillary changes. J Neuropathol Exp Neurol 58, 1010–1019.

Petry, F. R., Pelletier, J., Bretteville, A., Morin, F., Calon, F., Hebert, S. S., Whittington, R. A. and Planel, E. (2014) Specificity of anti-tau antibodies when analyzing mice models of Alzheimer’s disease: problems and solutions. PloS one 9, e94251.

Ren, Q. G., Liao, X. M., Chen, X. Q., Liu, G. P. and Wang, J. Z. (2007) Effects of tau phosphorylation on proteasome activity. FEBS Lett 581, 1521–1528.

Sankaranarayanan, S., Barten, D. M., Vana, L. et al. (2015) Passive immunization with phospho-tau antibodies reduces tau pathology and functional deficits in two distinct mouse tauopathy models. PloS one 10, e0125614.

Sereno, L., Coma, M., Rodriguez, M. et al. (2009) A novel GSK-3beta inhibitor reduces Alzheimer’s pathology and rescues neuronal loss in vivo. Neurobiol Dis 35, 359–367.

Steinhilb, M. L., Dias-Santagata, D., Fulga, T. A., Felch, D. L. and Feany, M. B. (2007) Tau phosphorylation sites work in concert to promote neurotoxicity in vivo. Molecular biology of the cell 18, 5060–5068.

Yanamandra, K., Kfoury, N., Jiang, H., Mahan, T. E., Ma, S., Maloney, S. E., Wozniak, D. F., Diamond, M. I. and Holtzman, D. M. (2013) Anti-tau antibodies that block tau aggregate seeding in vitro markedly decrease pathology and improve cognition in vivo. Neuron 80, 402–414.

Zheng-Fischhofer, Q., Biernat, J., Mandelkow, E. M., Illenberger, S., Godemann, R. and Mandelkow, E. (1998) Sequential phosphorylation of Tau by glycogen synthase kinase-3beta and protein kinase A at Thr212 and Ser214 generates the Alzheimer-specific epitope of antibody AT100 and requires a paired-helical-filament-like conformation. Eur J Biochem 252, 542–552.

