## Supplementary figures and images for "High specificity of widely used phospho-tau antibodies validated using a quantitative whole-cell based assay"

### Supplement Figures

##### Supplementary Figure 1.

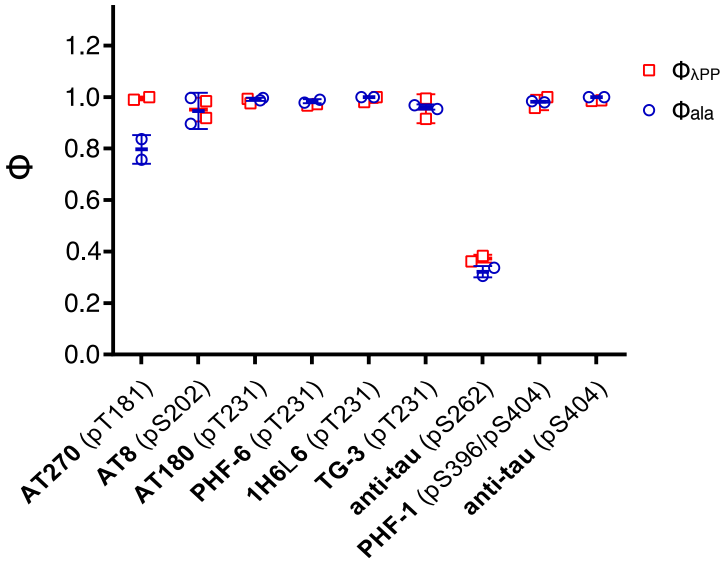

#### Supplementary Figure 2.

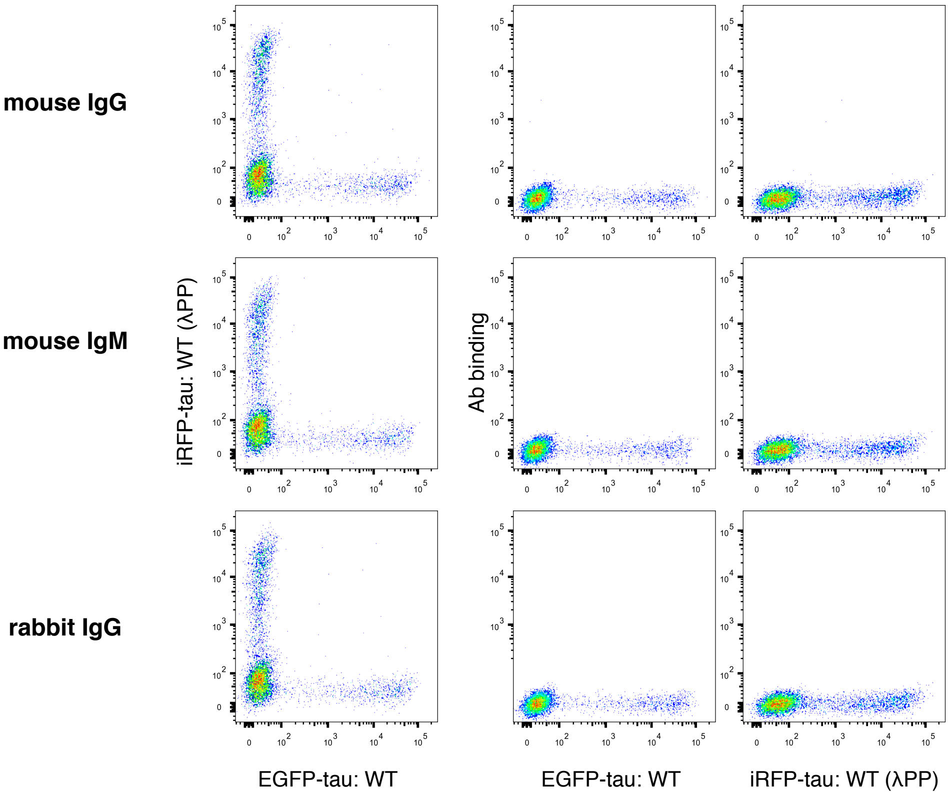
