## Supplement Tables for "High specificity of widely used phospho-tau antibodies validated using a quantitative whole-cell based assay"

***Supplementary Table 1.*** *Primers for Golden Gate assembly of plasmids*

| DNA template | Primers |
|---|---|
| pRK5-EGFP-Tau | 5'- TCCGCTGAGCCCCG -3'<br>5'- CCATGGTGGCGACCATC -3' |
| piRFP670-N1 | 5'- ATGGTAGCAGGTCATGCC -3'<br>5'- CGGATCCGCTCTCAAGCGCGG -3' |

***Supplementary Table 2.*** *Primers used for site-directed mutagenesis*

| Mutant | Primers |
|---|---|
| T181A | 5'- GAGCTGGGTGGTGCCTTTGGAGCGGGC -3'<br>5'- GCCCGCTCCAAAGGCACCACCCAGCTC -3' |
| S202A* | 5'- AGTGCCTGGGGCGCCGGGGCTGC -3'<br>5'- GCAGCCCCGGCGCCCCAGGCACT -3' |
| T212A/S214A | 5'- GTTGGAAGGGCCGGGGCGCGGGAGCGG -3'<br>5'- CCGCTCCCGCGCCCCGGCCCTTCCAAC -3' |
| T231A | 5'- GACTTGGGTGGAGCACGGACCACTGCCACCTTCT -3'<br>5'- AGAAGGTGGCAGTGGTCCGTGCTCCACCCAAGTC -3' |
| S262A | 5'- TTCAGGTTCTCAGTGGCGCCGATCTTGGACTTG -3'<br>5'- CAAGTCCAAGATCGGCGCCACTGAGAACCTGAA -3' |
| S396A/S404A§ | 5'- CAGACACCACTGGCGCCTTGTACACGATCTC -3'<br>5'- GAGATCGTGTACAAGGCGCCAGTGGTGTCTG -3' |
| S404A | 5'- AGATGCCGTGGAGCCGTGTCCCCAGAC -3'<br>5'- GTCTGGGGACACGGCTCCACGGCATCT -3' |

Note: *Previous study has shown that AT8 recognition requires only Ser202 to be phosphorylated in tau (Goedert *et al.* 1993).

§S396A/S404A were obtained using the S404A mutant as a template to generate double mutations.

**References**

Goedert, M., Jakes, R., Crowther, R. A., Six, J., Lubke, U., Vandermeeren, M., Cras, P., Trojanowski, J. Q. and Lee, V. M. (1993) The abnormal phosphorylation of tau protein at Ser-202 in Alzheimer disease recapitulates phosphorylation during development. *Proc Natl Acad Sci U S A* **90,** 5066-5070.
